# FuFiHLA: A tool for Full-Field HLA typing from long reads data

**DOI:** 10.1101/2025.10.23.684216

**Authors:** Jingqing Hu, Qian Qin, Heng Li, Ying Zhou

## Abstract

0.

**Motivation:** Allele typing for Human Leukocyte Antigen (HLA) genes has many important clinical applications. Popular short-read typing can only accurately distinguish alleles at the peptide level, which potentially limit our understanding of the effect of variants in non-coding region. Recently, a few methods were declared to distinguish full-field HLA alleles from de-novo assemblies, which motivates us to develop an accurate HLA typing method directly from long reads.

**Results:** We developed FuFiHLA, a lightweight open-source software, to type HLA alleles. Currently it supports typing alleles of six HLA genes (HLA-A, HLA-B, HLA-C, HLA-DRB1, HLA-DQA1, and HLA-DQB1) from long reads. Evaluation using 47 PacBio HiFi samples from HPRC shows that FuFiHLA achieves 99.57% accuracy in the full field allele typing and QV as 50.1 for consensus allele sequence construction. Additional testing on four Nanopore R10 reads demonstrates slightly reduced accuracy in the fourth field.

**Availability and implementation:** FuFiHLA is available on GitHub (https://github.com/jingqing-hu/FuFiHLA) under MIT License.

## 1. Introduction

The Human Leukocyte Antigen (HLA) region is located on the short arm of chromosome 6, specifically on band 6p21.3, and covers more than one hundred coding genes that play crucial roles in human immune system. This region has been implicated in over hundreds of diseases (1). Due to the need of recognizing unpredicted antigens, several HLA genes exhibit exceptionally high within-species diversity even within the coding region. For instance, pairwise differences are 3% for HLA-A, 4% for HLA-DRB1, and 5% for HLA-DQB1 (2), whereas the genome-wide average is 0.1% in human (3) and about 1.23% between human and chimpanzee (4).

Regarding the high polymorphism in HLA genes, a nomenclature system has been developed for distinguishing gene sequences or alleles (5). A specific gene sequence, or an allele, is assigned a name consisting of four fields, addressing the difference in antigen binding affinity (field 1), in peptide (field 2), in coding nucleotide (field 3), and in intron nucleotide (field 4). An extra suffix tag may be appended to denote expression changes.

HLA typing is an art of inferring the accurate HLA allele name from biospecimen based on a pre-exist reference database. In this work, we will focus on using PacBio HiFi reads of DNA sequences (6), and extending to Nanopore R10 data with slight modification. The IMGT/HLA is used as the standard reference data set for HLA allele typing (7). In the latest version 3.59, it consists of 41003 distinct alleles covering 47 HLA genes.

With a known reference allele data set, there are two major computational strategies for HLA allele typing from DNA sequences: one is to infer the best allele combinations from reference database to explain the observed reads, the other is to recover or reassemble the allele sequence and type the allele by comparing to the reference data set (8–12). Using short read data, the first strategy is less robust with undocumented alleles, while the second strategy might be affected by assembly errors. Thus, both strategies can hardly achieve high accuracy in full-field HLA typing.

Long read sequencing could be a game-changer for full field HLA typing. Because of the long extension of read length (about 15kb for PacBio HiFi read), it is much easier than using the short reads (less than 300bp) to recover full length sequence of HLA allele. Recently, several assembly-based methods claimed to have high typing accuracy in full field (2,13,14). In complementary, there is a timely need for long reads-based HLA typing method. So far there are three typing tools taking long reads but with limitations: HLA*LA(11) only outputs the first three fields nomenclature; Starphase (15) (a new version of HiFiHLA) only supports PacBio HiFi data; SpecImmune (16) needs an aligned bam with a specific version of reference genome, which is less convenient in practice. In this work, we will present a new method for HLA allele typing on the six graft transplant genes (HLA-A, HLA-B, HLA-C, HLA-DQA1, HLA-DQB1, and HLA-DRB1), it can take both aligned and raw/unaligned long reads as input, typing HLA allele in full-field and constructing consensus allele sequences. It supports both PacBio HiFi and Nanopore R10 data.

## 2. Pipeline Overview

This pipeline processes reads through sequential steps to output allele sequences and allele types. (Figure 1a) Alleles with only partial sequences in IMGT/HLA are ignored in our current typing.

**Figure 1.**
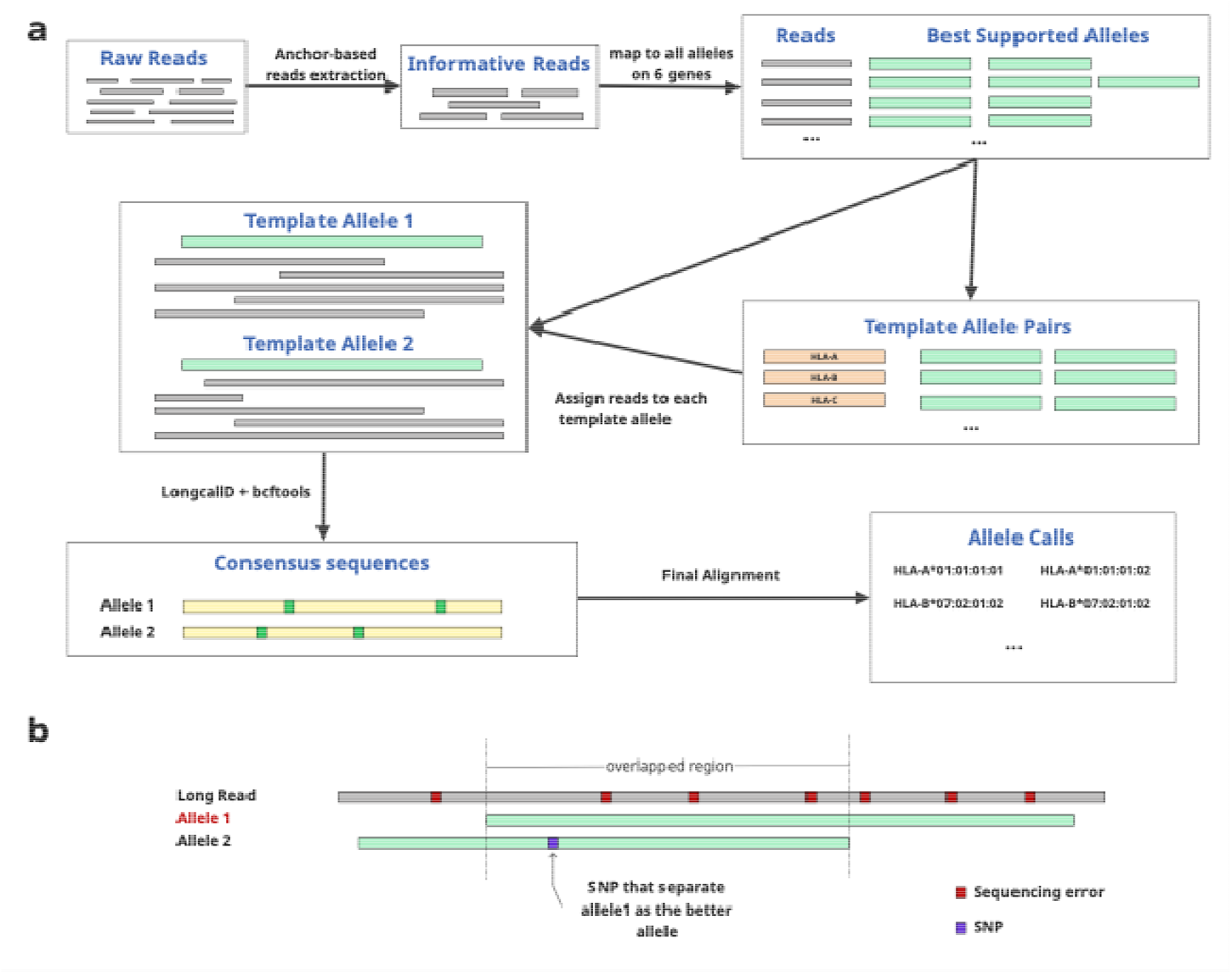
Overview of FuFiHLA. **a**) Informative reads are extracted from raw reads by mapping them to the anchor alleles and further mapped to all alleles from targeted genes to find out template allele pair for each gene. Then those informative reads are assigned to each of the template allele for variants calling and consensus gene sequence construction. Reconstructed sequences that include variants from template allele are further mapped to all alleles of targeted genes for allele typing. **b**) MOI method for choosing better alignment.

### 2.1 Informative Read extraction

We used a subset of IMGT/HLA alleles as anchors to extract informative reads from raw reads. When bam and the gene annotation of the reference are available, we can also use reads overlapped with the targeted genes as input. In our evaluation, both generated identical allele typing results, but with slightly difference in consensus sequence construction.

To construct anchor allele set, we clustered all IMGT/HLA allele sequences with cd-hit (identity cutoff = 0.95) (17), and from each cluster we selected one allele as anchor allele to represent that allele cluster. We also added allele sequences from HLA-DRB6, HLA-DRB7, HLA-DRB8, and HLA-DRB9, which are mostly partial, to the anchor set. In total there are 136 anchor alleles selected, covering the six targeted genes and 37 other HLA genes.

The reads were mapped to all anchor alleles and further extracted as informative reads based on the overlapping with anchor alleles from the six targeted genes. Particularly, an informative read should be covering at least 40% length of any targeted anchor allele or covering at least 200bps from one end of a targeted anchor allele. Due to the similarity among HLA genes, one read can be mapped to several alleles even from different genes which may introduce additional noise. Thus, the allele-to-read mappings were removed if the mismatch rate is 10 times higher than the lowest mismatch rate of the same read.

### 2.2 Alignment comparison

To determine the better alignment between two reads to an allele or two alleles to a read, we restrict the comparison of **M**ismatch in the **O**verlapped mapping **I**nterval (**MOI**) (Figure 1b). This design is tested to be more practical than using mismatch rate directly due to the existence of sequencing error and the nature of allele length variation. With MOI method, the better alignment includes a smaller number of mismatches in the overlapped mapping interval. For reads with higher error rates, such as the Nanopore R10 data, more ‘tie’ situations are created in the comparison by ignoring sequence difference with at most one indel or substitution event.

### 2.3 Template allele pair selection and consensus allele sequence construction

A pair of template alleles for each targeted gene are used as references to construct consensus allele sequences from informative reads, based on the assumption that the target sample has exact two copies of each of the six genes. Template allele pair is a combination of reference alleles of the highest agreement with the reads. In total 19,782 allele sequences of the targeted genes are used in the selection.

To select a pair of template alleles for each of the targeted genes in one sample, we constructed a pool of alleles (P_allele_) to iterate allele pairs and a pool of reads (P_read_) that belongs to the specific gene. When multiple alleles are mapped to the same region of a particular read, the best-match allele(s) are selected by the MOI method through pairwise comparison. The selected allele(s) are added to the allele pool P_allele_, and the supporting read is added to the read pool P_read_. The mappings between selected alleles and their supporting read are also recorded.

For an allele A, a three-metric tuple S(A) = (*n, c, m*) is calculated, where *n*is the total number of unique supporting reads, *c* is total read coverage, and *m* is total number of matched base pairs. If one read supports several alleles, then its contribution to *c* will be divided by the total number of supported alleles.

The three-metric tuple for a pair of alleles, noted A and B, is calculated as *S*(*A, B*) = *S*(*A*) + *S*(*B*). We rank allele pairs based on the three-metric tuple *S*(*A, B*) in decreasing order and the allele pair of the first rank is selected as the template allele pair. Practically, we found that top 15 alleles for HiFi reads and top 30 alleles for R10 reads with highest coverage was good enough to select the proper template allele pair.

Once the template allele pair is selected, reads are assigned/phased to each of them through MOI method. If the mappings to each template allele are equal, then the read is assigned to both alleles. Since it tends to give two different alleles even though for the homozygous sites, we added an additional filtration to the template alleles that if one allele’s mapping coverage is four times smaller than the other, the template allele with less coverage will be removed and the site is forced to be homozygous.

With phased reads and the template allele, we applied longcallD for variants calling (18) and “bcftools consensus” for allele sequence construction (19). Only major variants supported by at least three reads are used for variants calling and consensus sequence construction.

### 2.4 HLA Allele typing

HLA allele typing is based on a penalty score by considering different mismatch penalties of mapping between consensus allele sequence and reference allele sequence. Exonic mismatch penalty is 10 per event and intronic mismatch penalty is 1 per event. The allele with the lowest penalty score is selected as the typing allele.

## 3. Results

We first evaluated the performance of our HLA typing tool on 47 HiFi samples from the Human Pangenome Reference Consortium (Table S1), which includes 16 Americans, 24 Africans, 6 Asians and 1 European. The average read depth is 41X and average read length was about 15 kb. The error rate of HiFi long reads is <1%. (20)

Regarding the assembly-based annotation as ground truth (2), we compared FuFiHLA with three other long read typing tools, HLA*LA(1.0.4), Starphase(1.3.2) and SpecImmune(1.0.0), across the six targeted genes at different resolutions (Table 2). In our evaluation, both Starphase and FuFiHLA demonstrated high accuracy, exceeding 99% even in the full-field.

**Table 1.** Accuracy of long read HLA typing on six targeted genes.

| **Software** | **1-field** | **2-field** | **3-field** | **4-field** |
| --- | --- | --- | --- | --- |
| Starphase | 99.47% | 99.47% | 99.29% | 99.35% |
| HLA-LA | 98.94% | 85.79% | 79.04% | NA |
| SpecImmune | 96.99% | 89.34% | 86.50% | 71.82% |
| FuFiHLA (raw) | 100% | 99.82%* | 99.82%* | 99.57% |
| FuFiHLA (bam) | 100% | 99.82%* | 99.82%* | 99.57% |

\* 100% accuracy if ignoring the error caused by the coding sequence allele (HLA-DRB1\*04:92) on HG01358 that is not present in gene sequence database

**Table 2.** Runtime in minute. Alignment time for Starphase is the time used to construct bam file.

|  | sample | HG002 | HG00733 | HG01123 | NA19240 | HG00621 | HG03579 | HG02622 |
| --- | --- | --- | --- | --- | --- | --- | --- | --- |
| bam_size | (GB) | 33 | 78 | 91 | 98 | 111 | 122 | 134 |
| Alignment<br>(minute) | FuFiHLA (raw) | 270 | 772 | 889 | 941 | 919 | 1135 | 1060 |
|  | Starphase | 1759 | 4173 | 4889 | 5049 | 5674 | 7312 | 6899 |
|  | SpecImmune | 1883 | 4331 | 4907 | 5289 | 5874 | 7594 | 6901 |
|  | HLA*LA | 1759 | 4173 | 4889 | 5049 | 5674 | 7312 | 6899 |
| Typing<br>(minute) | FuFiHLA (raw) | 32 | 63 | 100 | 76 | 94 | 114 | 126 |
|  | FuFiHLA (bam) | 21 | 41 | 60 | 46 | 50 | 65 | 53 |
|  | Starphase | 3 | 4 | 4 | 3 | 3 | 5 | 4 |
|  | SpecImmune | 33 | 36 | 39 | 41 | 37 | 39 | 40 |
|  | HLA*LA | 255 | 714 | 572 | 682 | 588 | 1083 | 856 |

We also evaluate the base pair consistency of gene sequence construction by aligning the constructed gene sequences to the de novo assemblies of each of the 47 samples from HPRC phase 2. Among the 564 allele sequences, FuFiHLA (raw/unaligned read as input) gives 548 perfectly matches and 16 with mismatches. Among the 16 gene sequences with mismatched nucleotides, 12 of them have indels of homopolymer or short tandem repeats and four of them have single nucleotide substitutions. (Table S2) Using bam for reads extraction has similar performance. (Table S3) Meanwhile Starphase provided 532 perfect matches and 32 with mismatches. Among the 32 gene sequences with mismatched nucleotides, 26 of them have indels of homopolymer or short tandem repeats and five have single base substitutions and/or indel differences, and 1 wrong calling for DRB1 allele with edit distance larger than 4000.(Table S4) The overall QV score of consensus allele sequence is 50.1 for FuFiHLA and 44.6 for Starphase (one wrong call was excluded).

Using Nanopore R10 reads would lead to reduced accuracy for FuFiHLA in the fourth field. Among four testing samples (HG002 from GIAB (21) and three non-cancer samples from CASTLE panel (22)), it achieved 100% accuracy for the first three fields but dropped to 87.0% for the forth field, which mainly due to the sequencing error of homopolymers in introns.(Table S5)

## 4. Discussion

Long reads HLA typing is timely needed for clinical application. In this work we presented a new method, FuFiHLA, which can accurately type six HLA genes from WGS long reads. Based on the evaluation on 47 HPRC HiFi samples, FuFiHLA achieved 99.57% accuracy at 4-field allele typing. Starphase is another method developed recently by PacBio, which has similar typing accuracy as FuFiHLA but less accurate in consensus allele sequences construction.

Starpahse is faster than FuFiHLA when taking bam (aligned) as input, but slower when accounting for bam preparing time because using anchor sets is more efficient than constructing bam to extract informative reads. (Table 2)

Taking advantage of the large number of reference HLA gene sequences in IMGT/HLA, we are able to select template pair that minimizing the difference between reference alleles and targeted alleles which makes it more robust in allele typing and allele sequence construction, even though the reads depth is low. For example, the HG002 sample we used only has coverage of 10X but we can give all corrected allele typing for 12 alleles and only one allele with mismatches as indels of homopolymer or short tandem repeats.

One potential risk for estimation error could be caused by the in-completeness of IMGT/HLA reference data. The only allele typing mismatch in the second filed made by FuFiHLA in our evaluation happened on the allele DRB1*04:92 in the sample HG01358, where the allele’s full gene sequence is not available in the reference data and the coding sequence (partial) was used for typing the ground truth by Immuannot. However, FuFiHLA gave the identical allele sequence as the assembly.

In summary, we provided the most accurate method for full field HLA allele typing and consensus gene sequences construction on six graft transplant related genes. Currently it supports PacBio HiFi and Nanopore R10 reads, but the framework could be extended for covering more genes with other types of long reads sequencing data.

## Acknowledgements

We would like to acknowledge the National Genome Research Institute (NHGRI) for funding the following grants supporting the creation of the human pangenome reference: U41HG010972, U01HG010971, U01HG013760, U01HG013755, U01HG013748, U01HG013744, R01HG011274, and the Human Pangenome Reference Consortium (BioProject ID: PRJNA730823).

## Data availability

HPRC HiFi reads and assemblies are publicly available from https://humanpangenome.org/hprc-data-release-2/ and the list of 47 samples are included as supplementary file. Nanopore R10 reads include three non-cancer samples (H1437, H2009 and HCC1937) from CASTLE panel (https://github.com/CASTLE-Panel/castle) and HG002 from GIAB (https://42basepairs.com/browse/s3/ont-open-data/giab_2023.05/analysis/hg002/).

